# Intracellular Delivery of Nanoparticles *via* Microelectrophoresis Technique: Feasibility demonstration

**DOI:** 10.1101/2020.03.18.996173

**Authors:** Mengke Han, Jiangbo Zhao, Joseph Mahandas Fabian, Sanam Mustafa, Yinlan Ruan, Steven Wiederman, Heike Ebendorff-Heidepriem

**Affiliations:** Institute for Photonics and Advanced Sensing (IPAS) and School of Physical Sciences, The University of Adelaide, Adelaide, South Australia 5005, Australia; ARC Centre of Excellence for Nanoscale BioPhotonics (CNBP), The University of Adelaide, Adelaide, South Australia 5005, Australia; Adelaide Medical School, The University of Adelaide, Adelaide, South Australia 5005, Australia

**Keywords:** Intracellular delivery, Microelectrophoresis, Nanoparticles, Quantum dots, Biosensor

## Abstract

Nanoparticles with desirable properties and functions have been actively developed for various biomedical research, such as *in vivo* and *in vitro* sensors, imaging agents and delivery vehicles of therapeutics. However, an effective method to deliver nanoparticles into the intracellular environment is a major challenge and critical to many biological studies. Current techniques, such as intracellular uptake, electroporation and microinjection, each have their own set of benefits and associated limitations (*e.g.*, aggregation and endosomal degradation of nanoparticles, high cell mortality and low throughput). Here, the well-established microelectrophoresis technique is applied for the first time to deliver nanoparticles into target cells, which overcomes some of these delivery difficulties. Semiconductive quantum dots, with average hydrodynamic diameter of 24.4 nm, have been successfully ejected *via* small electrical currents (−0.2 nA) through fine-tipped glass micropipettes as an example, into living human embryonic kidney cells (roughly 20 - 30μm in length). As proposed by previous studies, micropipettes were fabricated to have an average tip inner diameter of 206 nm for ejection but less than 500 nm to minimize the cell membrane damage and cell distortion. In addition, delivered quantum dots were found to stay monodispersed within the cells for approximately one hour. We believe that microelectrophoresis technique may serve as a simple and general strategy for delivering a variety of nanoparticles intracellularly in various biological systems.

## INTRODUCTION

The noninvasive, intracellular delivery of exogenous materials with high efficiency and specificity, has shown great promise in deciphering and even modulating the complex, spatiotemporal interplay of biomolecules within living cells.^1–2^ Viral vectors, proteins, peptides, micelles, metabolites, membrane impermeable drugs, molecular probes and functionalized nanoparticles are all potential cargo materials for intracellular delivery.^1, 3^ For example, fluorescent semiconductive quantum dots (QDs) with superior optical properties and surface groups permit real-time tracking of intracellular molecules over time scales of milliseconds to hours, offering a capability to monitor intracellular events that cannot be accomplished *via* organic fluorophores.^2, 4–5^ In addition, QDs functionalized with specific receptor ligands have revealed the complex internalization mechanisms of growth factor receptors in endosomes, which advances the understanding of wound healing and embryonic development studies.^6–7^ With respect to gene therapy for treating severe inherited diseases, remedial genetic information has successfully passed through complex tissue and cellular barriers into target cells to replace the malfunctioning genes and proficiently expressed therapeutic molecules capable of reversing the condition.^1, 8^ Overall, the successful and efficient intracellular delivery of a broad range of exogenous cargo materials into diverse cell types is a critical pre-requirement to endow their specific biomedical functions.^1^

Current intracellular delivery strategies can be categorized into carrier-based and membrane-disruption-based modalities.^1^ Carrier-based techniques utilize the endocytosis or membrane-fusion pathway to release the carrier’s payload into living cells.^1, 3, 9^ The unique strengths of carrier-based techniques are the cell-type-specific uptake and subcellular-compartment-specific release, along with the appropriate spatiotemporal dynamics in response to stimuli, including alterations in pH, temperature, oxidation and light.^1, 10^ However, carrier systems are limited by feasible combinations of cargo materials and cell types.^1^ Cargo materials, with large variations in physiochemical properties (charge, size and functional groups), may not be efficiently packaged with the carriers, or unpacked properly (only about 1 - 2 % endosomal escape), leading to insufficient intracellular discharge.^1, 11^ Additionally, target cells often do not offer appropriate receptors to embrace the carriers or endosomal escape pathways to effectively discharge the cargoes.^1^

To alleviate the limitations of carrier-based strategies, membrane-disruption approaches, such as electroporation, microinjection and microelectrophoresis, have been widely studied and developed. In general, membrane-disruption approaches allow to physically introduce transient discontinuities in the plasma membrane for the intracellular delivery of cargo materials.^1, 5, 12^ The distinct advantages of these approaches are the high independence on the cargo properties or cell types as well as precisely controllable membrane-perturbing effects for instantaneous delivery.^1^

Among membrane-disruption approaches, microelectrophoresis technique uses small electrical currents to eject charged substances through fine-tipped glass micropipettes into target cells.^13^ Compared to microinjection and electroporation, which uses pressure and high-voltage electrical field impulse respectively,^2, 5, 12–13^ microelectrophoresis performs intracellular delivery in a more controlled manner for three reasons. Firstly, microelectrophoresis can limit the problematic diffusion of chemically and pharmacologically active substances from micropipettes, by simply applying a retaining current.^13^ This can reduce cell distortion and damage caused by the volume of solvent ejected *via* microinjection.^13^ Secondly, as most biological membranes *in vivo* maintain resting membrane potential differences ranging from −30 to −180 mV,^14^ microelectrophoresis can readily locate target cells, without the guidance of a microscope for microinjection.^15^ Once the micropipette is pierced into the cytosol of target cell, it can conveniently monitor its intracellular electrical activity in real-time.^16^ Thirdly, microelectrophoresis can minimize the physical damage to target cells by using small electrical currents rather than the high power impulses used in electroporation, which generate temporal pores in the cell membrane for transportation but can lead to high cell mortality.^2^

However, although microelectrophoresis technique has been established since *circa* 1900,^12^ no studies have been conducted on exploring the intracellular microelectrophoretic delivery of nanoparticles, despite the rapid development of utilizing nanomaterials in various intracellular biological research and medical applications.^2^ The main challenge confronting microelectrophoretic delivery of nanoparticles is the possibility of nanoparticle aggregation in the tip of micropipettes during ejection, which can cause tip blockage and failed delivery. The reasons are twofold.

Firstly, traditionally used silver/silver chloride (Ag/AgCl) electrodes in microelectrophoresis technique only conduct well (transform the flow of electrons from the current source to a flow of ions in solution) to locate target cells and subsequently record their intracellular electrical activity with high signal to noise ratio and wide recording bandwidth (only for electrically excitable cells, *i.e.*, neurons, muscle cells and some endocrine cells) in solutions that contain substantial Cl^−^ ions, *i.e.*, 2 - 4 M potassium chloride (KCl).^17^ For example, conventional fluorescent dyes for intra-cellular microelectrophoresis, such as biocytin and lucifer yellow are prepared in 2 M potassium acetate and 0.5 - 1 M lithium chloride.^16, 18^ However, for nanoparticles, this high concentration of KCl can significantly lower their repulsive energy barrier, *i.e.*, zeta potential at their hydrodynamic diameters (**Fig. S1**), making the adhesion of particles irreversible.^19^ This leads to blockages in the tip of micropipettes during ejection and failed microelectrophoresis. Therefore, it is important to prepare optimal nanoparticle suspensions with a suitable KCl concentration to maintain the colloidal stability of nanoparticles for ejection and at the same time permit high-fidelity intracellular recording.

Secondly, to impale cells with minimal damage, a rule of thumb is that the outer diameter (OD) near the tip of micropipettes should be 500 nm or less.^13^ However, the inner diameter (ID) near the tip must be large enough to allow the ejection of nanoparticles having comparable hydrodynamic diameters. Tips that are too small will impede the ejection and subsequently cause the aggregation of nanoparticles in the tips, leading to failed microelectrophoresis. Therefore, it is important to fabricate micropipettes with correct tip sizes. In parallel, the magnitude of ejecting current and duration must be tuned for optimal delivery.

In this paper, we demonstrate for the first time the proof of concept that microelectrophoresis is an effective and reproducible method for the intracellular delivery of nanoparticles by addressing the above related technical issues. We establish an efficient method for the preparation of nanoparticle suspensions that balance the colloidal stability against the intracellular recording quality. We describe the fabrication of micropipettes with correct tip sizes and successful intracellular microelectrophoresis with appropriate ejecting current and duration. These results suggest the extensive potential of microelectrophoresis technique as a simple and precise approach in the intracellular delivery of various nanoparticles in the future.

## RESULTS AND DISCUSSION

### Optimizing nanoparticle suspension for intracellular delivery

In this study, cadmium selenide/zinc sulfide (CdSe/ZnS) core-shell structured QDs (emission maxima of 655 nm) with amine-derivatized polyethylene glycol (PEG) surface functional group (Invitrogen, US), hereafter referred to as 655-QDs, were used to demonstrate intracellular microelectrophoresis.

Successful microelectrophoretic delivery and intra-cellular recording require concentrated KCl solutions in micropipettes to attain sufficient conductivity, whereas the high KCl concentration can weaken the colloidal stability of nanoparticles. Thus, the colloidal stability of 655-QDs was investigated in the presence of different KCl concentrations using zeta potential and particle size distribution measurements, to find the optimal preparation of 655-QDs suspensions for successful microelectrophoresis and intracellular recording.

Zetasizer Nano ZSP (Malvern Instruments, UK) with a 633 nm red laser was used to measure the zeta potential of 655-QDs in aqueous media *via* laser Doppler electrophoresis.^20^ Particles with zeta potential more positive than 30 mV or more negative than −30 mV are generally considered to represent sufficient repulsion to maintain their colloidal stability.^20^ It is worth noting that sufficient conductivity of the suspending medium is required for reliable zeta potential measurements, so that the electrical field can be applied across the folded capillary cells without inducing electrode polarization and voltage irregularities.^21^ For zeta potential measurements in 0 M KCl solution, the original 655-QDs suspension in 50 mM borate buffer from the supplier is pipetted into 1.5 mL fresh ultrapure water, which leads to an extremely low borate concentration of 6.25e-5 M and thus a poor average conductivity of 1.9e-4 S/m (n = 6), which is close to the conductivity of ultrapure water, 5.5e-6 S/m as specified by the vendor Merck Millipore. In addition, when ultrapure water is exposed in an air atmosphere upon dispensing, it immediately absorbs carbon dioxide (CO_2_) and forms carbonic acid, which can cause a decrease in the pH with susceptibility to temperature fluctuations.^22^ These factors can lead to great variations in the measured zeta potential values of different 655-QDs samples in 0 M KCl.

The Zetasizer was also used to analyze the size distribution of 655-QDs in aqueous media *via* dynamic light scattering (DLS). DLS measures the time-dependent fluctuation of scattered light intensity caused by the constant Brownian motion of particles, and reports their hydrodynamic diameters as the equivalent hydrodynamic diameters (***D***_***H***_) of spheres that have the same average diffusion coefficient.^23^ Note that in ultrapure water at pH 7, the thickness of the electrical double layer of all particles is considered to be about 1 μm (**Fig. S**1),^24^ making nanoscale particle size distribution measurement in solution *via* DLS impossible. An established criterion for monodispersed nanoparticles is that their hydrodynamic diameters (***D***_***H***_) should be less than twice of their diameters in the dry state (***D***_***T***_) measured by transmission electron microscope (TEM).^25^ **Fig. 1a** shows the image of 655-QDs (dark dots) on the surface of a TEM grid. The average shape of 655-QDs is modelled as a prolate ellipsoid with the major axis (***a***_***T***_) of 9.7 ± 1.6 nm and the minor axis (***b***_***T***_) of 6.7 ± 0.8 nm (± 1 standard deviation (SD), n = 82) rather than ideal spheres. Therefore, as per the criterion for nanoparticle monodispersity in aqueous environment, monodispersed 655-QDs should theoretically have major hydrodynamic axes (***a***_***H***_) in the range of 8.1 nm - 22.6 nm and minor hydrodynamic axes (***b***_***H***_) in the range of 5.9 nm - 15.0 nm, which are less than twice of their ***a***_***T***_ and ***b***_***T***_ in the dry state. To compare with the results reported by DLS measurements, the following equation regarding the diffusion properties of anisotropic particles in Brownian motion,^26^ was used to translate the ellipsoidal dimensions (***a***_***H***_ and ***b***_***H***_) of 655-QDs to an equivalent diameter (***D***_***H***_) of spheres having the same diffusion coefficient:

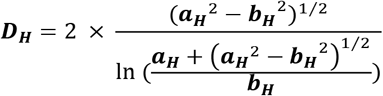

**Figure 1.**
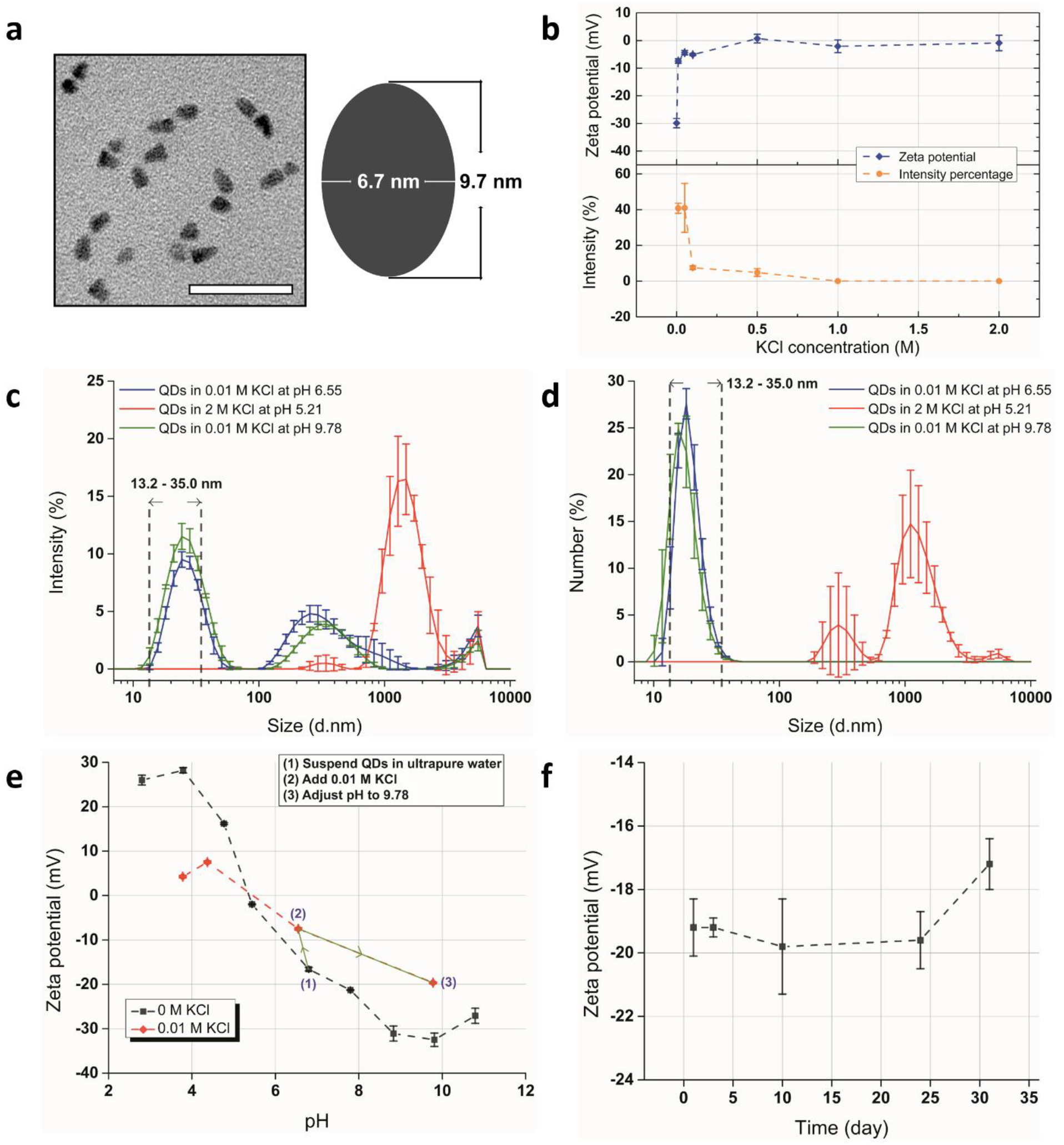
655-QDs suspension preparation. **a,** TEM image of 655-QDs reveals an average shape of prolate ellipsoid with a major axis (***a***_***T***_) of 9.7 ± 1.6 nm and a minor axis (***b***_***T***_) of 6.7 ± 0.8 nm (± 1 SD with n = 82). Scale bar, 50 nm. **b,** the zeta potential of 655-QDs and the fractions of light intensity scattered by monodispersed 655-QDs as a function of KCl concentration. Error bars, ± 1 SD with n = 3. **c,** the size distribution by intensity and **d,** by number of 655-QDs in 0.01 M (pH 6.55), 2 M KCl (pH 5.21) and optimized suspensions (0.01 M KCl adjusted to pH 9.78). Error bars, ± 1 SD with n = 3. The dot lines indicate the size range of monodispersed 655-QDs from 13.2 to 35.0 nm. **e,** the zeta potential of 655-QDs in ultrapure water and 0.01 M KCl solution with different pH values. Error bars, ± 1 SD with n = 3. Inserted with each step of the optimal preparation process of 655-QDs suspension for microelectrophoresis. **f,** the stability of 655-QDs zeta potential in optimized suspension. Error bars, ± 1 SD with n = 3.

In view of the range of ***a***_***H***_ and ***b***_***H***_ dimensions, mono-dispersed 655-QDs are considered to have hydrodynamic diameters ***D***_***H***_ over 13.2 nm and less than 35.0 nm.

Zeta potential measurements show that 655-QDs exhibit negative surface charge in ultrapure water, *i.e.*, 0 M KCl, leading to an average zeta potential of −29.9mV (**Fig. 1b**, upper panel, n = 3), which agrees with a previous report on the zeta potential of gold nanoparticles that are also surface-functionalized with amine-derivatized PEG.^27^ As the KCl concentration increases, the zeta potential (colloidal stability of 655-QDs) rapidly approaches zero due to the stronger electrostatic screening effect,^19^ which is accompanied by an irreversible aggregation of QDs as quantified by DLS measurements.

The fraction of light intensity scattered by monodispersed 655-QDs sharply decreases from 40.8 % in 0.01 M KCl to 7.5 % in 0.1 M KCl (**Fig 1b**, bottom panel, n = 3, no data on ultrapure water as noted before). **Fig. 1c** reveals the scattered light intensity of particles across a range of sizes in 0.01 M and 2 M KCl solutions. The dotted lines indicate the size range of monodispersed 655-QDs from 13.2 to 35.0 nm. In 0.01 M KCl, 59.2 % of the scattered light comes from QDs aggregates or artefacts (*e.g.*, dust). However, scattered light intensity is proportional to the sixth power of the particle radius and therefore the intensity-based size distribution is highly sensitive to very small numbers of aggregates or dust.^28^ Thus, the number of QDs aggregates is negligible compared to the total number of particles in the sample (determined using Mie theory),^28^ as shown in **Fig. 1d**. In 2 M KCl, QDs completely aggregate with a mean size around 1.5 μm due to the stronger electrostatic screening effect caused by the high electrolyte concentration.^19^

Considering the zeta potential and size distribution of 655-QDs in different KCl solutions, 0.01 M is an efficient KCl concentration for maintaining their colloidal stability. However, the zeta potential of −7.4 mV for 655-QDs in 0.01 M KCl solution is still not sufficiently low. Thus, to further stabilize 655-QDs, the relationship between the pH of suspension and the zeta potential of 655-QDs was investigated (**Fig. 1e**). The zeta potential in ultrapure water (*i.e.*, 0 M KCl) was measured to be − 16.6 mV. Note that this value is different to the prior study of KCl effect (−29.9 mV in **Fig. 1b**, upper panel) due to low conductivity of the suspension and large uncertainty of zeta potential measurements in ultrapure water as mentioned above.

The addition of alkali (NaOH) to the media results in a more negative charge for 655-QDs particles (decreased zeta potential). Conversely, the addition of acid (HCl) increases the zeta potential. The maximal zeta potential (absolute value) of −32.5 mV is at around pH 9.81 where the most stable state of 655-QDs exists. Although a stable state of 655-QDs also exists at around pH 3.79, a strong acid environment (pH<4) is not recommended by the supplier, where their polymer coating can dissociate, exposing and dissolving the core-shell structure. In addition, due to the high mobility of hydrogen ions (H^+^), a large amount of H^+^ in microelectrophoresis can result in lowering of the pH in the vicinity of the tip of micropipettes.^29^ This localized change in pH has been proposed to excite the cell undergoing intracellular recording and interfere with the normal physiological state.^30^ On the contrary, 655-QDs do not degrade in a strong basic environment (pH>9) as noted by the supplier. Furthermore, in comparison to the electrophoretic mobility of H^+^ (36.25 μmcm/Vs water at 25.0 ° C), hydroxide ion (OH^−^) has a lower electrophoretic mobility (20.50 μmcm/Vs in water at 25.0 °C), resulting in less effect on the intracellular activity.^31^

**Fig. 1e** reveals that pH adjustment can also buffer the negative effect of KCl on the stability of 655-QDs. With the same KCl concentration of 0.01 M in suspension, 655-QDs have a larger zeta potential (absolute value) at pH 3.78, 4.37 and 9.78 (zeta potential of −18.2 mV), than that at pH 6.55 without any pH adjustment. Therefore, the optimal protocol for the preparation of 655-QDs suspension for microelectrophoresis is to initially disperse QDs in ultrapure water and then add 2 M KCl to the suspension until a final concentration of 0.01 M achieved. Finally, the pH should be adjusted to 9.78 to further stabilize QDs. The green curve in **Fig. 1c** shows the size distribution of optimized 655-QDs suspension, where 53.9 % of scattered light comes from monodispersed QDs that constitute 91.4 % of the total number of particles in the sample as **Fig. 1d** shows.

For practical microelectrophoresis applications, preparation of fresh suspensions would be too time-consuming. A stock suspension with good colloidal stability and ready for use would be highly beneficial. **Fig. 1f** shows the shelf life of optimized 655-QDs suspensions (0.01 M KCl at pH 9.78), which were aliquoted and stored in several 1.5 mL eppendorf tubes sealed with parafilm in a 4.0 °C refrigerator. The zeta potential values of QDs in these intact aliquots were measured on different days, which remained the same for at least 24 days, indicative of this required long-term colloidal stability.

### The effect of KCl concentration on the quality of intracellular recording

The KCl concentration of 0.01 M has been demonstrated to be sufficient to maintain the colloidal stability of 655-QDs. The next step is to test whether such a low electrolyte concentration, suitable for nanoparticles in suspension, allows for the recording of intracellular activity with high fidelity in real-time. Here, we compared the responses captured from visual neurons, binocular small target motion detector (BSTMD2), in the optic lobes of two dragonflies.^32^ One used micropipettes filled with 2 M KCl (used in standard dragonfly electrophysiology), and the other filled with optimized 655-QDs suspension (0.01 M KCl at pH 9.78). Micropipettes were inserted with a working Ag/AgCl electrode from the blunt end and held by a micromanipulator to slowly approach target neurons (**Fig. 2a**). Aluminosilicate capillaries were used to fabricate micropipettes with program 1 using a horizontal micropipette puller (P-97; Sutter Instrument, US), resulting in extremely fine tips (*ca.* 100 nm tip ID, measured in previous studies of standard dragonfly electrophysiology)^33^ to impale dragonfly neurons with negligible damage (**Fig. 3a**, upper panel, **Table S**1). Their resistances were measured by applying a small current, *e.g.*, 1 nA, through the micropipette from the bridge amplifier to confirm that there was no blockage or breakage in the tips. A decrease in resistance implies breakage of the tip, whereas an increase indicates that the tip is blocked. No sign has occurred during recording. Micropipette filled with 2 M KCl solution has tip resistance of 120 MΩ on average and 335 MΩ on average for optimized 65-QDs suspension, respectively.

**Figure 2.**
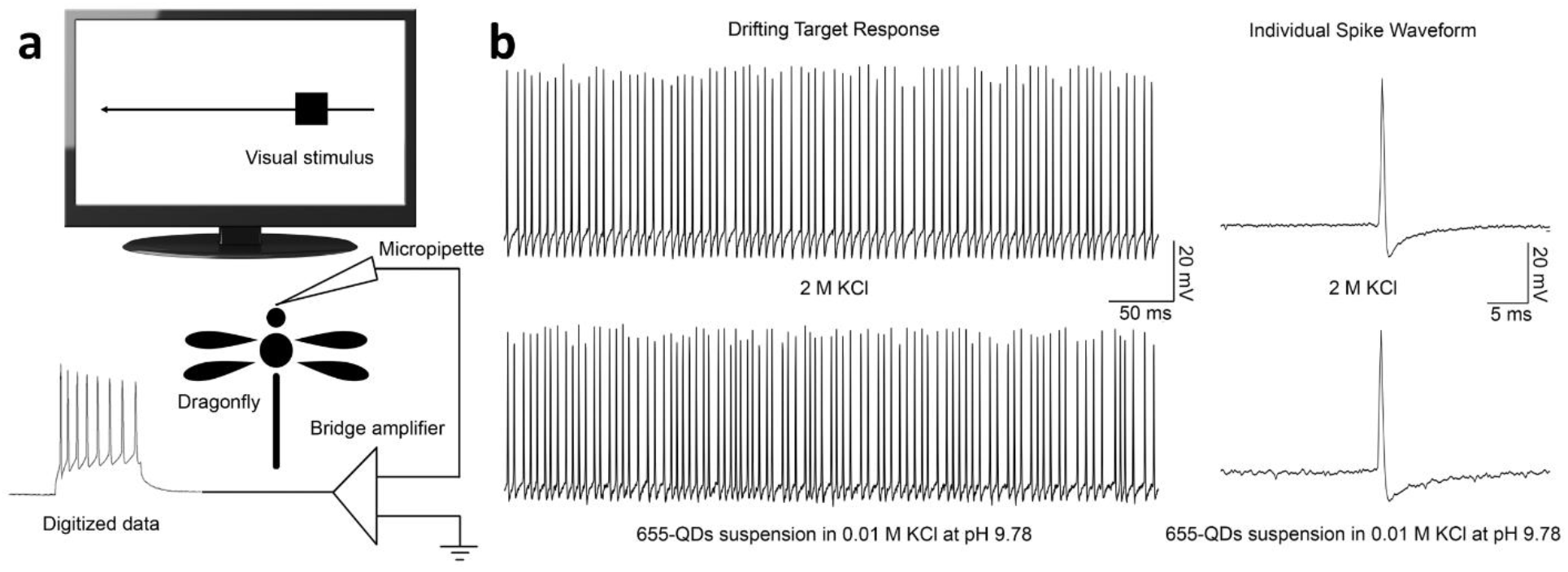
The effect of KCl concentration on the quality of intracellular recording. **a,** schematic illustration of experiment setup for intracellular recording of dragonflies. A liquid crystal display (LCD) monitor was placed in front of the dragonfly for stimulating visual neurons by drifting small moving objects. The visual stimulus elicited voltage changes across the cell membranes of single lobula neurons, which were recorded in real-time. **b,** the responses of two BSTMD2 cells in two separate dragonflies to the presentation of a drifting object, which were recorded with micropipettes filled with 2 M KCl solution and optimized 655-QDs suspension (0.01 M KCl at pH 9.78), respectively.

**Figure 3.**
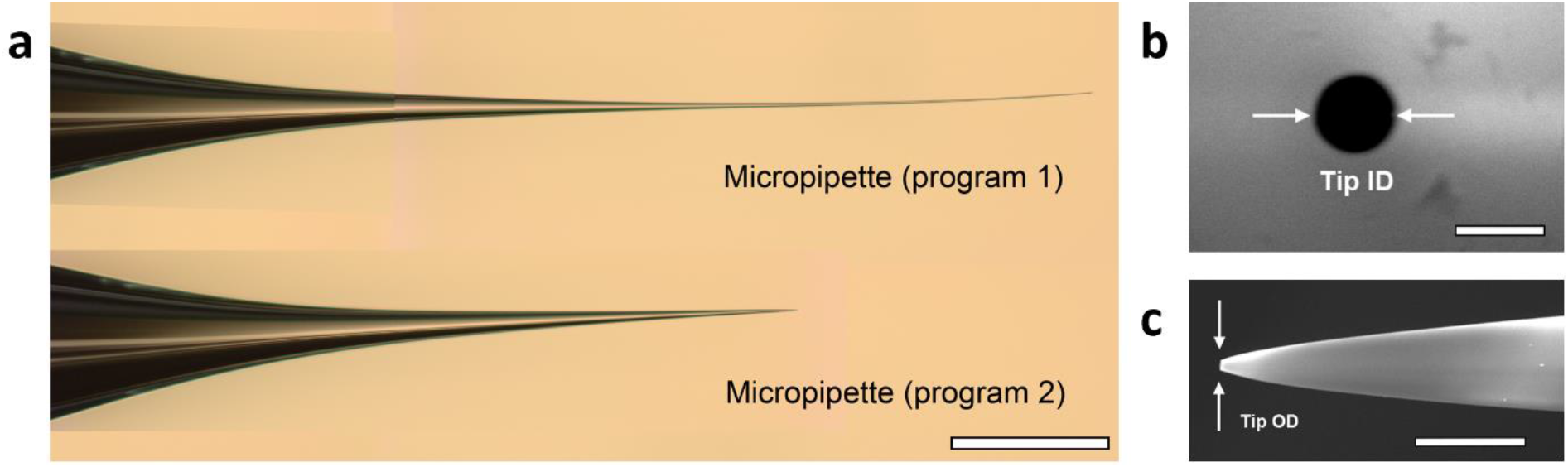
Micropipette fabrication. **a,** optical microscope image of two micropipettes fabricated with pulling program 1 for intracellular electrophysiological recordings of dragonfly neurons (upper panel) and pulling program 2 for microelectrophoresis of 655-QDs (bottom panel), respectively. Scale bar, 500 μm. **b,** high resolution SEM image of a micropipette for microelectrophoresis of 655-QDs (pulled with program 2 in **Table S**1) with a tip ID of 210.7 nm (front view). The orifice of micropipette is the black circle near the centre of the image. Scale bar, 250 nm. **c,** high resolution SEM image of another micropipette (pulled with program 2) with a tip OD of 212.2 nm (side view). Scale bar, 2.5 μm.

When BSTMD2 is presented with a small drifting target, the cell responds by significantly increasing the frequency of action potential firing. **Fig. 2b** shows the raw responses (left panel) and an enlarged view of individual spike waveforms (right panel) from the two separate BSTMD2 cells presented with a small moving target. Spiking responses remained very similar for both electrolyte concentrations. In addition, the individual action potential waveforms recorded by optimized 655-QDs suspension filled electrodes show only minor increases in noise. The same recording procedure was performed in several target-detecting neurons in the dragonfly. The micropipettes filled with 0.01 M KCl solution (suitable for 655-QDs suspension) had an average tip resistance of 300 ± 85 MΩ (±1 SD, n = 6). In these recordings, we observed a greater degree of variation in quality (i.e., noise and signal amplitude). However, in all cases, it was feasible to sort spikes that were sufficiently distinct from the resting potential without any issue in temporal responsiveness.

As a conclusion, KCl concentration of 0.01 M in suspensions is efficient to maintain the colloidal stability of 655-QDs and precisely locate target cells, following with high-fidelity intracellular recordings.

### Optimizing the tip size of micropipette for intracellular delivery

For a successful microelectrophoresis, the tip ID of the micropipette is required to be larger than the sum of hydrodynamic diameters of nanoparticles and other dissolved ions that pass through the tip for conductivity. The range of hydrodynamic diameter of monodispersed 655-QDs is 13.2 - 35.0 nm. The theoretical hydrated diameters of K^+^, Cl^−^ and Na^+^ ions are 0.3, 0.4 and 0.2 nm, respectively.^34^ Therefore, the tip ID of micropipette should be over 50 nm to allow the ejection, but the tip OD should be less than 500 nm to avoid physical damage to target cells.^13^

In this study, the IDs and ODs of aluminosilicate capillaries and the tip IDs and ODs of micropipettes fabricated with pulling program 2 (**Fig. 3a**, bottom panel, **Table S**1) using P-97 micropipette puller for microelectrophoresis were measured by an optical microscope (**Fig. S**2) and a scanning electron microscope (SEM) (**Fig. 3b** and c, and **Fig. S**3), respectively. Compared to typically used borosilicate glass, aluminosilicate glass provides increased hardness, improved chemical durability and reduced electrical conductivity.^35^ The ratio of tip ID to tip OD in a borosilicate micropipette is consistent over its total taper length, whereas in aluminosilicate micropipettes, the ratio increases remarkably towards the tip,^35^ which allows extremely fine tips to be formed and satisfies the requirement for successful intracellular microelectrophoresis.

The ID and OD of aluminosilicate capillaries were measured to be 0.52 mm and 0.99 mm on average for 26 capillaries, with variations of ± 0.03 and ± 0.02 mm (± 1 SD), respectively, which are within the vendor’s specification of ± 0.05 mm. When pulling micropipettes, capillaries with slightly different IDs and ODs will have different distances to the box heating filament and different volume of air enclosed in the internal channel, which can alter the glass temperature and result in variations in tip ID and OD of micropipettes.^36^ In addition, this effect is enlarged due to the inherent fluctuations of heating temperature, air pressure and moisture, and capillary emplacement in the puller, though model P-97 is designed with good reproducibility. Moreover, the reliability of each puller is different and largely dependent on its fine settings.

To measure the tip IDs and ODs with high accuracy, the fabricated micropipettes were fixed in two different orientations on the SEM sample holder; either vertically for tip IDs or horizontally for tip ODs measurement (**Fig. S**3b and c). Thus, it was not possible to measure both the ID and the OD for the same micropipette tip. The average tip OD is 202 nm with a tolerance of ± 35 nm (± 1 SD, n = 26). The average tip ID of micropipettes is 206 nm with a larger tolerance of ± 46 nm (± 1 SD, n = 26), which is partially caused by the observational error due to the inconsistency of pipette angle when manually fixing micropipettes onto the vertical SEM sample holder. These two averages are nearly identical, which validates the unique characteristics of aluminosilicate glass to enable the fabrication of extremely fine tips with greatly increased tip ID to OD ratio of ~1.0 compared to ID/OD ratio of 0.5 for the capillaries. In summary, the range of tip IDs of micropipettes is suitable for the ejection of 655-QDs and the tip ODs avoid the physical damage to target cells simultaneously.

### Successful intracellular delivery of nanoparticles into living cells *via* microelectrophoresis

The diagram of microelectrophoresis experiment is shown in **Fig. 4a**. Human embryonic kidney (HEK293) cells at roughly 20 - 30 μm in length,^37^ were seeded at 80,000 cells/well onto a chambered slide and cultured for one day prior to the microelectrophoresis of 655-QDs to achieve a confluency of 80 %. The micropipette filled with optimized QDs suspension and inserted with the working Ag/AgCl electrode from the blunt end was held by a micromanipulator and slowly approached single cells at the bottom of the well. A change in potential difference to around −38 mV indicated that the tip of micropipette was successfully pierced into the cytoplasm of HEK cells.^37^ Following, a small current of −0.2 nA was applied to eject QDs into the cell for 3 minutes. Since the cell confluency is 80 %, the tip of micropipettes will have a 20 percent chance of hitting the blank bottom surface of the culture slide where no cell is attached. Thus, the resistance of micropipettes was frequently measured to confirm that there was no blockage or breakage in the tips. If any of the two signs occurred, the micropipette was replaced. We didn’t observe any tip blockage sign during microelectrophoresis, which demonstrates an ideal monodispersity of optimized 655-QDs suspension and proper choice of tip ID for ejection. The resistance of several micropipettes pulled by program 2 varied from 50 MΩ to 80 MΩ due to the variation in their tip sizes and remained the same when removed out of the cells after delivery. Finally, the slide was placed in a stage top incubator and observed under a confocal laser scanning microscope to confirm the successful intracellular delivery of QDs.

**Figure 4.**
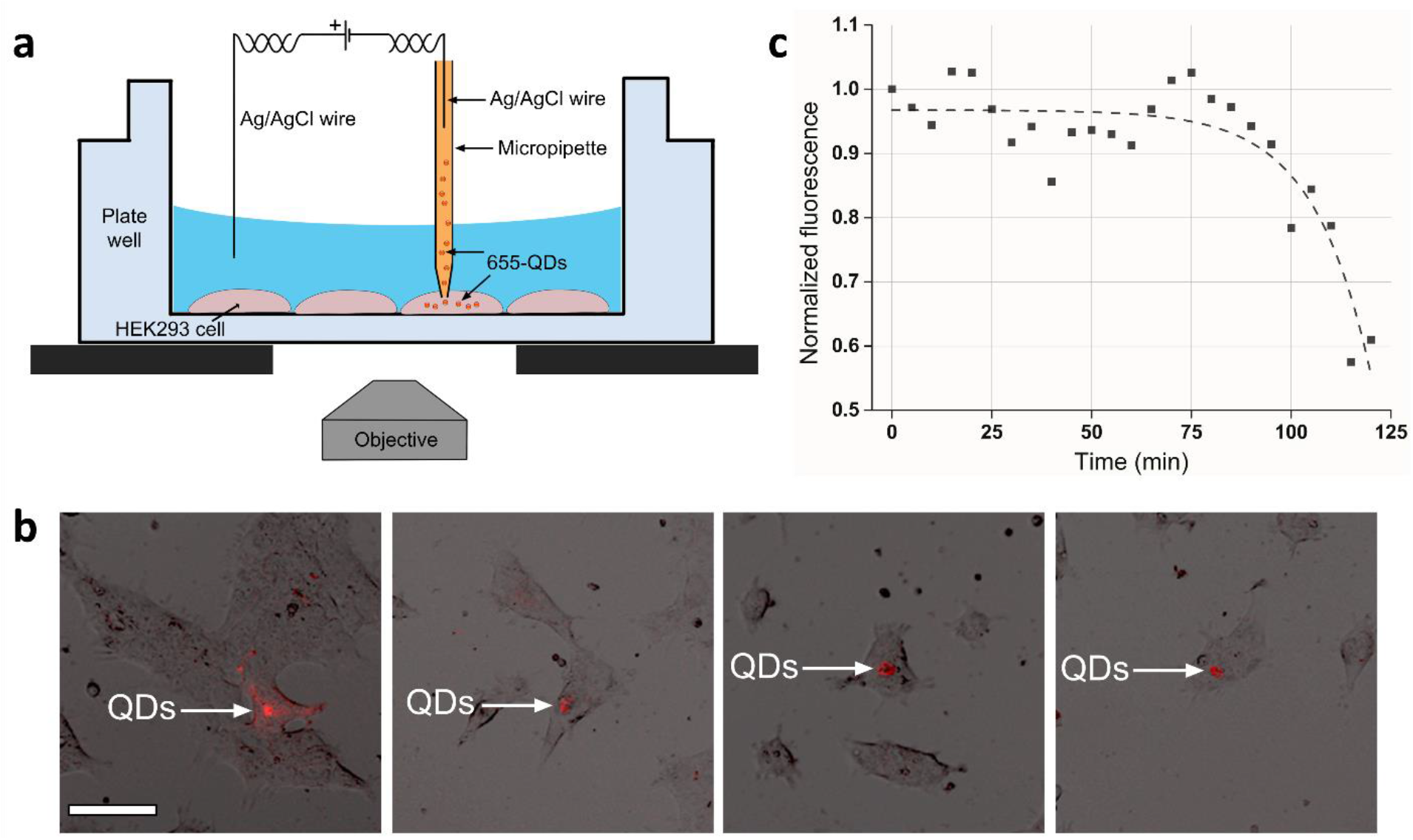
Microelectrophoresis of 655-QDs into living HEK293 cells. **a,** diagram of microelectrophoresis of 655-QDs into HEK cells and the position of objective in confocal imaging. **b,** overlay images of HEK cells with microelectrophoretic-delivered 655-QDs. The red dots in the cytosol are 655-QDs with arrows indicated. Scale bar, 10 μm. **c,** the fluorescence intensity changes of delivered 655-QDs in a HEK cell as a function of time. All data points are fitted to a dashed curve using the one-phase exponential decay function, *I* = *I*_0_ − 9.4*e*^*x*/14.3^, with R^2 = 0.79, where *I*_0_ denotes the initial intensity.

**Fig. 4b** shows the overlay images of four HEK293 cells delivered with 655-QDs *via* microelectrophoresis. QDs are present primarily in the cytosol and spread from the injection point to the whole cell. The amount of delivered QDs varies among cells, which results from the variation in tip ID of micropipettes. However, the calculation of the amount of delivered 655-QDs is not feasible due to the complicated determination and measurement of transport number in Faraday’s laws.^12^ The movement of delivered 655-QDs was then monitored for 2 hours (5 minutes/capture, **Movie S**1, Supporting Information). **Fig. 4c** shows the normalized intensity change of the fluorescence from delivered QDs in a living HEK cell in these 2 hours. The fluorescence intensity of 655-QDs shows no measurable loss for approximately one hour but then gradually decreases, which might be caused by the exocytosis or fluorescence quenching due to slow aggregation in the cellular environment.^38^ The aggregation of 655-QDs can be caused by unsuitable pH values of subcellular compartments, from 4.7 in lysosome to 8.0 in mitochondria,^39^ and high intracellular ionic strength of K^+^ at 139 mM,^40^ as shown in **Fig. 1b** and e.

## CONCLUSIONS

In summary, we demonstrate for the first time that the well-established microelectrophoresis technique can deliver nanoparticles intracellularly, such as QDs used here, and permit high-quality intracellular recordings at the same time. This was achieved by overcoming the following three critical challenges. Firstly, we prepared QDs suspensions with high colloidal stability to prevent aggregation and blockages in the tip of micropipettes during ejection. Secondly, we recorded the intracellular electrical activity of dragonfly neurons with high fidelity using micropipettes filled with optimized QDs suspensions. Thirdly, we fabricated micropipettes with tip sizes large enough to allow the ejection of QDs and as small as possible to avoid physical damage to target HEK293 cells.

This successful microelectrophoretic ejection of QDs lays the foundation for further studies and applications of microelectrophoresis technique for the intracellular delivery of various nanoparticles, which have been used as intracellular sensors, deep tissue and tumor imaging agents, and vectors for studying nanoparticle-mediated drug delivery.^41^ Our approach overcomes some of the limitations of current techniques, such as the endosomal degradation of nanoparticles in carrier-based strategy, high cell mortality and aggregation of nanoparticles in electroporation, and the cell locating difficulties in microinjection. In contrast, micro-electrophoresis technique can easily locate the position of target cells and precisely deliver monodispersed nanoparticles into target cells with negligible cell membrane damage and cell distortion. Delivered nanoparticles were found to stay monodispersed within the cells for approximately one hour. In addition, it can record the intracellular electrical activities of target cells at the same time, thus indicating pathological effects of the injected materials.

However, microelectrophoresis can lead to plasma membrane discontinuities that are slightly inconsistent from cell to cell caused by the variations in tip sizes of micropipettes fabricated with the same pulling program. The result may be either insufficient or excessive delivery that leads to cell damage. In addition, as microelectrophoresis is highly dependent on the membrane perturbation, it is restricted to adherent cells. Future studies should focus on minimizing the variation of the ID and OD of capillaries and improving the reproducibility of pipette pullers to fabricate micropipettes with less tip size variations, which is critical to the reproducibility of microelectrophoresis and cell membrane damage. Furthermore, fluorescent dyes, such as propidium iodide, can be used to evaluate and confirm cell viability after microelectrophoresis to identify how much damage is caused to the cells. Moreover, it is worth studying the volume of nanoparticle suspension and its impact on the activity of the cells and intracellular pH value in the future.

## EXPERIMENTAL SECTION

### The effect of KCl concentration and pH on the colloidal stability of nanoparticles

To vary the concentration of KCl, the original 8 μM 655-QDs suspension in borate buffer (50 mM borate) from the supplier was pipetted into 1.5 mL fresh ultrapure water (Merck Millipore, US) of varying concentrations of KCl (Chem-supply, AU). To vary the pH of QDs suspensions, the original 655-QDs suspension was firstly pipetted into 1.5 mL fresh ultrapure water and then the suspension pH was adjusted by gradually adding 0.1 M Hydrochloric Acid (HCl) or 0.1 M Sodium Hydroxide (NaOH) (Chem-supply, AU). For each sample, the suspension pH and temperature readings were recorded until stable (change in pH value was less than ±0.05 unit within 1 minute). To demonstrate that the pH adjustment can buffer the negative effect of KCl on the stability of QDs, the original 655-QDs suspension was pipetted into fresh ultrapure water of 0.01 M KCl. Then the suspension pH was adjusted by gradually adding 0.1 M HCl or 0.1 M NaOH to certain values.

All operations were conducted in a clean fume cupboard. The pH meter (Metrohm, CH) was calibrated by three standard pH buffers (VWR, US) at pH 4.00 ± 0.02, 7.00 ± 0.02 and 9.22 ± 0.02 at 20.0 °C. The original 655-QDs suspension was gently vortexed for 1 minute before the dilution. The concentration of 655-QDs was consistently 10 nM in each sample. KCl, HCl and NaOH solutions were centrifuged at 4000 revolutions per minute (rpm) for 1 minute before the addition to remove any large-size impurities that can affect measurement results.

Zetasizer nano ZSP was used for the studies on the colloidal stability of 655-QDs and was capable of both hydrodynamic size analysis and zeta potential measurements. Samples were pipetted into 1.5 mL Eppendorf Flex-tubes respectively and sonicated (Soniclean, AU) from 4.0 °C to 24.0 °C without the use of external heat for 20 minutes to obtain a better dispersion. Then, the samples were pipetted by 1 mL syringes into clean folded capillary cells (DTS1070; Malvern Instruments, UK) for subsequent measurements in Zetasizer.

In Zetasizer, Henry’s function was set at the value of 1.50.^42^ The dispersant was set to be water (Temperature: 25.0 °C; Viscosity: 0.8872 cP; Refractive Index: 1.330; Dielectric constant: 78.5) and its viscosity was used as the viscosity of the sample. The refractive index and absorption of 655-QDs were set as 2.550 and 0.010.^43^ The temperature equilibrium time was set as 120 seconds at 25.0 °C. The real-time temperature value was recorded when pH reading was carried out since the measurement temperature in Zetasizer can cause fluctuations of actual pH values during measurement. But, the change of pH value of each sample is the same for the same temperature and has no influence on data interpretation.

### Intracellular recording of dragonfly neuron

Wild-caught dragonflies (*Hemicordulia tau*) were immobilized with a mixture of beeswax and gum rosin (solid form of resin) (1:1) on a plastic articulating stage as shown in **Fig. 2a**. To gain the access to the brain surface, a small hole was dissected on the posterior surface of the head capsule. A working Ag/AgCl electrode (782500; A-M Systems, US) was connected to an intracellular bridge mode amplifier (BA-03X; npi electronic, DE) and a counter Ag/AgCl electrode was inserted into the head capsule surface to form a complete electrical circuit. With a pipette holder (PPH-1P-BNC; ALA Scientific Instruments, US) and a micromanipulator (MM-33; ALA Scientific Instruments, US), micropipettes (pulled with program 1, **Table S**1) were pierced into single lobula neurons. Neurons were stimulated by drifting small moving visual features across a high refresh rate (165 Hz) LCD monitor placed directly in front of the dragonfly. Data were digitized at 5 kHz with a 16-bit analog-to-digital (A/D) converter and analyzed off-line with MATLAB. The visual stimulus elicited voltage changes across the cell membranes of neurons and the digitized data indicated the successful intracellular recording of neurons in real time.

### Micropipette fabrication for microelectrophoresis of nanoparticles

A horizontal micropipette puller, model P-97 Flaming/Brown (Sutter Instrument, US) and aluminosilicate glass capillaries (OD = 1.0 mm, ID = 0.53 mm, Length = 100 mm) with filaments (Harvard Apparatus, US) were used in this study to fabricate micropipettes with correct tip sizes for microelectrophoresis of 655-QDs.

Capillaries were carefully fixed onto a horizontal metal stage and imaged under an optical microscope (BX51; Olympus, JP) to measure their IDs and ODs (**Fig. S**2). The parameter settings of the pulling program 2 are presented in **Table S1** in the Supporting Information. Fabricated micropipettes were carefully fixed onto carbon tape covered metal stages (**Fig. S**3) and coated with a 3 nm-thick platinum film for imaging under a FEI Quanta 450 FEG environmental SEM to measure their tip IDs and ODs.

### Cell culture

HEK293 cells were seeded at 80,000 cells/well onto slides (μ-Slide 8 well with chambered coverslip; ibidi; DE) and incubated at 37 °C in 5 % CO_2_ overnight in 300μL Dulbecco’s modified Eagle’s media (DMEM) (Sigma-Aldrich, US) supplemented with 2 mM L-glutamine (Sigma-Aldrich, US) and 10 % fetal bovine serum (Gibco, US). The next day, when the cells approached a confluency of 80 % on the slide, the media were replaced by pre-warmed 250 μ-L Hank’s balanced salt solution (HBSS) supplemented with 25 mM 4-(2-hydroxyethyl)-1-piperazineéthanesulfonic acid (HEPES, no phenol red) just prior to the microelectrophoresis.

### Microelectrophoresis of nanoparticles into living cells

#### Nanoparticle suspension preparation

Firstly, the original 655-QDs suspension was pipetted into fresh ultrapure water inside a glass beaker (Corning Incorporated, US) with a magnetic stir bar rotating continuously at approximately 200 rpm, resulting in a QDs concentration of 10 nM. Then, 2 M centrifuged KCl solution was gradually added into the diluted QDs suspension to a final KCl concentration of 0.01 M. Finally, 0.1 M centrifuged NaOH solution was gradually added into the suspension to reach a stable pH value at 9.78. The optimized QDs suspension was aliquoted into several 1.5 mL Eppendorf Flex-tubes and sealed with parafilm and stored in a 4 °C refrigerator. The colloidal stability of QDs as a function of time was investigated by measuring the zeta potential of one intact aliquot on the different days.

Before microelectrophoresis, a stored 655-QDs suspension aliquot was gently vortexed for 1 minute and sonicated from 4.0 °C to 24 °C without the use of external heat for 20 minutes to obtain a better dispersion. Then it was centrifuged at 3000 rpm for 40 seconds to remove any large-size impurities and aggregates to avoid tip blockage during ejection. Finally, the supernatant (around 1 mL) was pipetted by a 1 mL syringe and carefully backfilled into clean micropipettes *via* a flexible plastic syringe needle (Warner instruments, CT).

#### Microelectrophoresis

The diagram of experiment apparatus is shown in **Fig. 4a**. The cell culture slide was placed on a vibration-proof table in a dark environment to avoid the photo-bleaching of 655-QDs. The micropipette filled with 655-QDs suspension and inserted with a Ag/AgCl working electrode from the blunt end was held by the pipette holder on the micromanipulator. The micropipette was slowly inserted into the cell culture medium *via* the micromanipulator. Another Ag/AgCl counter electrode was carefully placed into the medium. The two electrodes were connected to the headstage of the intra-cellular bridge mode amplifier formed a complete electrical circuit. The tip was slowly pierced into single HEK cells when a potential difference was recorded. Then a-0.2 nA current was applied to eject 655-QDs into HEK cells for 3 minutes.

### Confocal laser scanning microscope

A confocal laser scanning microscope (FV3000; Olympus, JP) equipped with a stage top incubator (Tokai Hit, JP) was used to obtain the overlay images (405 nm excitation) of living HEK293 cells that were delivered with 655-QDs *via* microelectrophoresis and track the movement and fluorescence intensity changes of these QDs over time.

## ASSOCIATED CONTENT

### Supporting Information

The Supporting Information is available on the website at DOI: xxx.

The following sections are included: Zeta potential and hydrodynamic diameter, the fabrication and size measurement of micropipettes (PDF) Movie S1: The fluorescence intensity changes of microelectrophoresis-delivered 655-QDs in living HEK cells for 2 hours

## AUTHOR INFORMATION

### Corresponding Author

*

### Notes

All authors have given approval to the final version of the manuscript. The authors declare no competing fi-nancial interest.

## Supporting information

The movement of delivered 655-QDs was then monitored for 2 hours (5 minutes/capture, Movie S1, Supporting In-formation).

## ACKNOWLEDGEMENTS

This work was performed in part at the Optofab node of the Australian National Fabrication Facility (ANFF) utilizing Commonwealth and SA State Government funding. The authors acknowledge partial support from the Australian Research Council Centre of Excellence for Nanoscale BioPhotonics (CNBP) (CE140100003) and Discovery Early Career Researcher Award (DECRA) from Australian Research Council (ARC) (DE150100548). M.H. thanks Ken N., Lyn W. and Agatha L. and Jane S. for their assistance in SEM, TEM, fluorescence and confocal laser scanning images acquisition at Adelaide Microscopy, and Jingxian Y. for useful discussion.

## REFERENCES

1. Stewart, M. P.; Sharei, A.; Ding, X.; Sahay, G.; Langer, R.; Jensen, K. F., In Vitro and Ex Vivo Strategies for Intracellular Delivery. Nature 2016, 538 (7624), 183–192.

2. Chou, L. Y.; Ming, K.; Chan, W. C., Strategies for the Intracellular Delivery of Nanoparticles. Chem. Soc. Rev. 2011, 40 (1), 233–245.

3. Stewart, M. P.; Lorenz, A.; Dahlman, J.; Sahay, G., Challenges in Carrier-Mediated Intracellular Delivery: Moving Beyond Endosomal Barriers. Wiley Interdiscip. Rev.: Nanomed. Nanobiotechnol. 2016, 8 (3), 465–478.

4. Pinaud, F.; Clarke, S.; Sittner, A.; Dahan, M., Probing Cellular Events, One Quantum Dot at a Time. Nat. Methods 2010, 7, 275–285.

5. Sun, C.; Cao, Z.; Wu, M.; Lu, C., Intracellular Tracking of Single Native Molecules with Electroporation-Delivered Quantum Dots. Anal. Chem. 2014, 86 (22), 11403–11409.

6. Lidke, D. S.; Lidke, K. A.; Rieger, B.; Jovin, T. M.; Arndt-Jovin, D. J., Reaching out for Signals: Filopodia Sense Egf and Respond by Directed Retrograde Transport of Activated Receptors. J. Cell Biol. 2005, 170 (4), 619–626.

7. Lidke, D.; Nagy, P.; Heintzmann, R.; Arndt-Jovin, D.; Post, J.; Grecco, H.; A Jares-Erijman, E.; M Jovin, T., Quantum Dot Ligands Provide New Insights into Erbb/Her Receptor-Mediated Signal Transduction. Nat. Biotechnol. 2004, 22, 198–203.

8. Naldini, L., Gene Therapy Returns to Centre Stage. Nature 2015, 526 (7573), 351–360.

9. Sahay, G.; Alakhova, D. Y.; Kabanov, A. V., Endocytosis of Nanomedicines. J. Controlled Release 2010, 145 (3), 182–195.

10. Scarpa, E.; Bailey, J. L.; Janeczek, A. A.; Stumpf, P. S.; Johnston, A. H.; Oreffo, R. O.; Woo, Y. L.; Cheong, Y. C.; Evans, N. D.; Newman, T. A., Quantification of Intracellular Payload Release from Polymersome Nanoparticles. Sci. Rep. 2016, 6, 29460.

11. Gilleron, J.; Querbes, W.; Zeigerer, A.; Borodovsky, A.; Marsico, G.; Schubert, U.; Manygoats, K.; Seifert, S.; Andree, C.; Stöter, M., Image-Based Analysis of Lipid Nanoparticle-Mediated Sirna Delivery, Intracellular Trafficking and Endosomal Escape. Nat. Biotechnol. 2013, 31 (7), 638–646.

12. Lalley, P. M., Microiontophoresis and Pressure Ejection. In Modern Techniques in Neuroscience Research, Windhorst, U.; Johansson, H., Eds. Springer Berlin Heidelberg: Berlin, Heidelberg, 1999; pp 193–212.

13. Curtis, D. R., Microelectrophoresis. New York: Academic Press: 1964; Vol. 5, p 144–190.

14. Tekle, E.; Astumian, R. D.; Chock, P. B., Electro-Permeabilization of Cell Membranes: Effect of the Resting Membrane Potential. Biochem. Biophys. Res. Commun. 1990, 172 (1), 282–287.

15. Chou, L. Y.; Ming, K.; Chan, W. C., Strategies for the Intracellular Delivery of Nanoparticles. Chem. Soc. Rev. 2011, 40 (1), 233–245.

16. Mobbs, P.; Becker, D.; Williamson, R.; Bate, M.; Warner, A. In Techniques for Dye Injection and Cell Labelling, Microelectrode techniques. The Plymouth workshop handbook. Cambridge, UK: The Company of Biologists Ltd, 1994; pp 361–387.

17. Axon Instruments, I., The Axon Guide for Electrophysiology & Biophysics Laboratory Techniques. Third ed.; Axon Instruments: 1993; p 117–118.

18. Hanani, M., Lucifer Yellow – an Angel Rather Than the Devil. J. Cell. Mol. Med. 2012, 16 (1), 22–31.

19. Zhang, W., Nanoparticle Aggregation: Principles and Modeling. In Nanomaterial: Advances in Experimental Medicine and Biology, 2014/04/01 ed.; 2014; Vol. 811, pp 19–43.

20. Clogston, J. D.; Patri, A. K., Zeta Potential Measurement. In Characterization of Nanoparticles Intended for Drug Delivery, McNeil, S. E., Ed. Humana Press: Totowa, NJ, 2011; pp 63–70.

21. Caputo, F., Measuring Zeta Potential. EUNCL-PCC-002 2015.

22. Corporation, Y., Understanding Ultrapure Water and the Difficulties with Ph Measurement. 2015.

23. Pecora, R., Dynamic Light Scattering: Applications of Photon Correlation Spectroscopy. Springer Science & Business Media: 2013; p 251.

24. Israelachvili, J. N., Intermolecular and Surface Forces. Academic press: 2011; p 313.

25. Moon, J.; Choi, K.-S.; Kim, B.; Yoon, K.-H.; Seong, T.-Y.; Woo, K., Aggregation-Free Process for Functional Cdse/Cds Core/Shell Quantum Dots. J. Phys. Chem. C 2009, 113 (17), 7114–7119.

26. Perrin, F., Mouvement Brownien D’un Ellipsoide (Ii). Rotation Libre Et Dépolarisation Des Fluorescences. Translation Et Diffusion De Molécules Ellipsoidales. J. Phys. Radium 1936, 7 (1), 1–11.

27. Xia, X.; Yang, M.; Wang, Y.; Zheng, Y.; Li, Q.; Chen, J.; Xia, Y., Quantifying the Coverage Density of Poly(Ethylene Glycol) Chains on the Surface of Gold Nanostructures. ACS Nano 2012, 6 (1), 512–522.

28. Mie, G., Beiträge Zur Optik Trüber Medien, Speziell Kolloidaler Metallösungen. Ann. Phys. (Berlin, Ger.) 1908, 330 (3), 377–445.

29. Gruol, D. L.; Barker, J. L.; Huang, L. Y.; MacDonald, J. F.; Smith, T. G., Jr., Hydrogen Ions Have Multiple Effects on the Excitability of Cultured Mammalian Neurons. Brain Res 1980, 183 (1), 247–252.

30. Frederickson, R. C.; Jordan, L. M.; Phillis, J. W., The Action of Noradrenaline on Cortical Neurons: Effects of Ph. Brain Res 1971, 35 (2), 556–560.

31. Duso, A. B.; Chen, D. D. Y., Proton and Hydroxide Ion Mobility in Capillary Electrophoresis. Anal. Chem. 2002, 74 (13), 2938–2942.

32. O’Carroll, D., Feature-Detecting Neurons in Dragonflies. Nature 1993, 362 (6420), 541–543.

33. O’Carroll, D. C. In *Biomimetic Visual Detection Based on Insect Neurobiology*, Electronics and Structures for MEMS II, International Society for Optics and Photonics: 2001; pp 1–11.

34. Marcus, Y., Ionic Radii in Aqueous Solutions. Chem. Rev. 1988, 88 (8), 1475–1498.

35. Instrument, S., P-97 Pipette Cookbook. 2008; p 65.

36. Chen, M. J.; Stokes, Y. M.; Buchak, P.; Crowdy, D. G.; Foo, H. T.; Dowler, A.; Ebendorff-Heidepriem, H., Drawing Tubular Fibres: Experiments Versus Mathematical Modelling. Opt. Mater. Express 2016, 6 (1), 166–180.

37. Thomas, P.; Smart, T. G., Hek293 Cell Line: A Vehicle for the Expression of Recombinant Proteins. J. Pharmacol. Toxicol. Methods 2005, 51 (3), 187–200.

38. Noh, M.; Kim, T.; Lee, H.; Kim, C.-K.; Joo, S.-W.; Lee, K., Fluorescence Quenching Caused by Aggregation of Water-Soluble Cdse Quantum Dots. Colloids Surf., A 2010, 359 (1), 39–44.

39. Casey, J. R.; Grinstein, S.; Orlowski, J., Sensors and Regulators of Intracellular Ph. Nat. Rev. Mol. Cell Biol. 2010, 11 (1), 50–61.

40. Lodish, H.; Berk, A.; Zipursky, S. L.; Matsudaira, P.; Baltimore, D.; Darnell, J., Intracellular Ion Environment and Membrane Electric Potential. In Molecular Cell Biology. 4th Edition, WH Freeman: New York, 2000.

41. Delehanty, J. B.; Mattoussi, H.; Medintz, I. L., Delivering Quantum Dots into Cells: Strategies, Progress and Remaining Issues. Anal Bioanal Chem 2009, 393 (4), 1091–1105.

42. Henry, D. In The Cataphoresis of Suspended Particles. Part I. The Equation of Cataphoresis, Proceedings of the Royal Society of London A: Mathematical, Physical and Engineering Sciences, The Royal Society: 1931; pp 106–129.

43. Hondow, N.; Brydson, R.; Wang, P.; Holton, M. D.; Brown, M. R.; Rees, P.; Summers, H. D.; Brown, A., Quantitative Characterization of Nanoparticle Agglomeration within Biological Media. J. Nanopart. Res. 2012, 14 (7), 1–15.

