## Supplemental Data 1 for "Intracellular Delivery of Nanoparticles *via* Microelectrophoresis Technique: Feasibility demonstration"

### S1. Zeta potential and hydrodynamic diameter

In colloidal dispersions, the net charge on the particle surface attracts oppositely charged ions in the surrounding interfacial region close to the surface, forming an electrical double layer structure (**Fig. S**1).[2, 3] The slipping plane is a notional boundary within the diffuse layer to divide ions, which forms a stable entity and moves with the particle at the same speed.[4] The diameter of the particle and the thickness of the slipping plane make the total hydrodynamic diameter.[4] The potential existing at the edge of the hydrodynamic diameter is the zeta potential.[4]

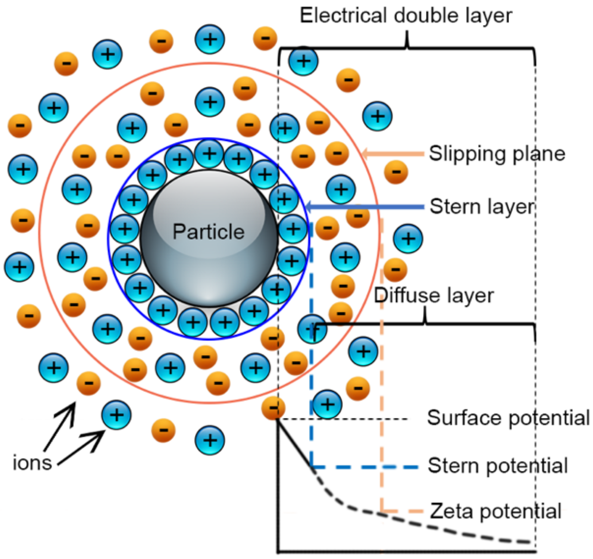


**Figure S1 | Schematic illustration of electrical double layer structure and zeta potential:** potential difference as a function of distance from the charged surface of a particle suspended in a medium. Reproduced with permission from reference 3. Copyright 2010 The Royal Society.

### S2. The fabrication and size measurement of micropipettes

**Table S1** The parameters of pulling program 1 and 2 in P-97 micropipette puller.

| Program | | 1 | 2 |
|---|---|---|---|
| Ramp | | 525 | 525 |
| Pressure | | 525 | 500 |
| Cycle 1 | Heat | 520 | 520 |
| | Pull | 0 | 0 |
| | Velocity | 8 | 8 |
| | Time | 1 | 1 |
| Cycle 2 | Heat | 515 | 475 |
| | Pull | 180 | 180 |
| | Velocity | 70 | 70 |
| | Time | 100 | 100 |

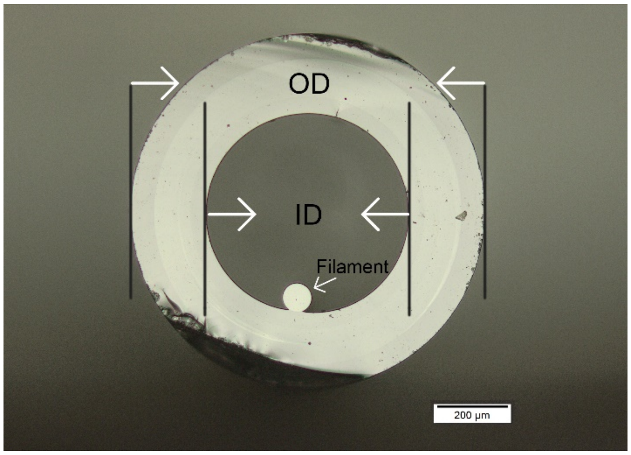


**Figure S2 | Optical microscope image of the cross section of an aluminosilicate glass capillary.** The inner diameter and outer diameter of the capillary are indicated with white arrows and black lines, respectively. The small sphere attached to the inner wall of the glass capillary is a glass rod referred to as filament with approximately 160 μm in diameter, to guide solutions from the blunt end to the tip of fabricated micropipettes *via* capillary force.[1] Scale bar, 200 μm.

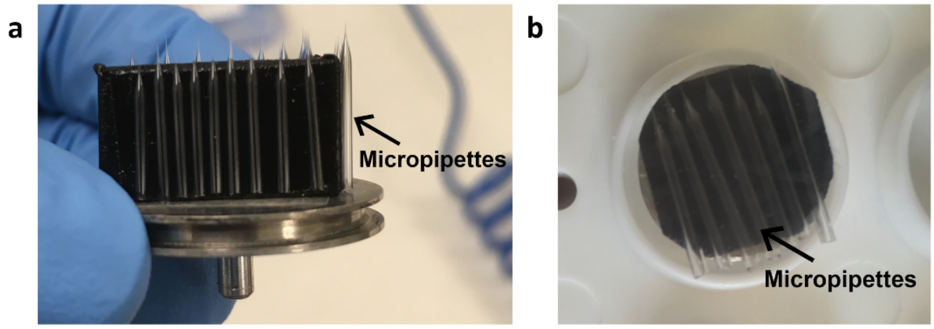


**Figure S3 | Placement of micropipettes on metal sample holders for SEM imaging. a,** micropipettes are fixed onto a vertical metal stage covered with carbon tape for the measurement of tip IDs. **b,** micropipettes are fixed onto a horizontal metal stage covered with carbon tape for the measurement of tip ODs.

### References


1. A. Oesterle, *P-97 Pipette Cookbook*, Instrument, Sutter, Novato, CA, 2008.
2. W. Zhang, in *Nanomaterial: Advances in Experimental Medicine and Biology*, 2014/04/01 edn., 2014, vol. 811, pp. 19-43.
3. M. Kaszuba, J. Corbett, F. M. Watson and A. Jones, *Philos. Trans. R. Soc., A*, 2010, **368**, 4439-4451.
4. A. Collins, *Nanotechnology cookbook: practical, reliable and jargon-free experimental procedures*, Elsevier, 2012.
